# Repeated SARS-CoV-2 Antigenic Exposures from Prior Vaccinations and Infections Demonstrate Limits of Antibody Durability and Breadth Against Newer Variants

**DOI:** 10.64898/2026.04.15.718804

**Authors:** Wei Wang, Emilie Goguet, Sabrina Lusvarghi, Stephanie Paz, Leeza Shrestha, Russell Vassell, Simon Pollett, Edward Mitre, Carol D. Weiss

**Affiliations:** Center for Biologics Evaluation and Research, US Food and Drug Administration, Silver Spring, Maryland; Department of Microbiology and Immunology, Uniformed Services University of the Health Sciences, Bethesda, Maryland; Department of Preventive Medicine and Biostatistics, Infectious Disease Clinical Research Program, Uniformed Services University of the Health Sciences, Bethesda, Maryland; Henry M. Jackson Foundation for the Advancement of Military Medicine, Inc., Bethesda, Maryland

**Keywords:** SARS-CoV-2, COVID-19 vaccines, neutralizing antibodies, XBB.1.5, JN.1, BA.3.2, antibody durability, immunological imprinting

## Abstract

**Background:** Widespread immunity from vaccination and infection has reduced COVID-19 morbidity and mortality, but this immunity varies across the population. Understanding how repeated antigenic exposures influence antibody responses helps to inform future vaccination strategies.

**Methods:** Serum samples collected one and six months after XBB.1.5 vaccination from 25 generally healthy healthcare workers with varying exposure histories were assessed for neutralizing activity against a range of variants, from pre-Omicron variants to latest Omicron JN.1 sublineage variants and divergent BA.3.2 variants, using lentiviral pseudoviruses. Participants were stratified by vaccination and infection history.

**Results:** XBB.1.5 vaccination elicited broad neutralizing responses, with strong boosting against previously encountered antigens relative to vaccine-matched XBB.1.5 and newer variants. Geometric mean neutralization titers were generally comparable across exposure groups, indicating limited influence of prior Omicron infection or bivalent vaccination, though intra-group heterogeneity was observed. At six months, overall titers declined by 36-62%. Titers remained highest against the pre-Omicron and lowest against JN.1 sublineage variants. N-terminal glycosylation (DelS31, T22N) modestly affected neutralization.

**Conclusions:** XBB.1.5 vaccination elicited broad neutralizing antibody responses against previously encountered and vaccine-matched antigens regardless of exposure history, but titers waned after six months. This waning, compounded by continued emergence of immune-evasive variants and heterogenous population immunity, underscores the need for continually monitoring neutralizing antibody durability and breadth to guide evidence-based COVID-19 vaccine formulation updates.

## BACKGROUND

The continuous evolution of SARS-CoV-2 has generated variants with increasing antigenic diversity. Early Omicron variants, including BA.1, BA.2, and BA.5, demonstrated significant immune evasion, a trend continued by XBB and JN.1 sublineages [1-4]. The recent emergence of BA.3.2, with over 40 mutations distinct from JN.1, highlights the ongoing evolution of SARS-CoV-2 that may evade existing immunity [5].

COVID-19 vaccine antigen formulations have been periodically updated to improve immune responses against new, antigenically divergent variants. Because rapid viral evolution prevents timely clinical trials, neutralizing antibody titers, which have been associated with protection [6], have been used to guide these updates. Spike mutations that increase neutralization escape often signal the need for such updates.

Periodic vaccine doses also help counter waning immunity [7, 8]. Studies have consistently demonstrated that neutralizing antibody (nAb) levels peak shortly after exposure, decline over the next six months, and then decline more slowly or stabilize at a lower level [9, 10]. Repeat antigenic exposures, however, lead to enhanced antibody affinity and memory responses over time [11-13]. Age, immune status, and prior SARS-CoV-2 infections influence the magnitude and kinetics of antibody response [14-16]. Hybrid immunity from a combination of vaccinations and infections has been associated with more durable and broader antibody responses [17-19]. However, the long-term durability of these responses after repeated antigenic exposures in the face of new variants remains poorly characterized.

As SARS-CoV-2 has become endemic, population immunity has become increasingly heterogeneous due to diverse combinations of antigenic exposures from vaccinations and infections [20]. Declining uptake of updated vaccines has further contributed to waning immunity and a more complex immune landscape. Such heterogeneity complicates predictions about the persistence and breadth of population immunity, which is important information for guiding vaccine updates and public health communication, especially when uptake of COVID-19 vaccine among high-risk populations remains low.

Here, we longitudinally analyzed neutralizing antibodies in multiply immunized healthcare workers with well-documented vaccine and infection histories, one and six months after receiving the 2023-2024 XBB.1.5 vaccine. We assessed neutralization against historical, vaccine-matched, late Omicron JN.1, KP.3, KP.3.1.1, XEC, LF.7, LP.8.1.1, NB.1.8.1, XFC and XFG variants, and the divergent BA.3.2 variants. These data provide insights into how repeated and varied antigenic exposures influence the durability and breadth of humoral immunity, informing evidence-based updates to COVID-19 vaccine formulation.

## MATERIALS AND METHODS

### Ethics statement

The Uniformed Services University Institutional Review Board approved the Prospective Assessment of SARS-CoV-2 Seroconversion (PASS) study (Protocol IDCRP-126), including details of the inclusion and exclusion criteria published before [21]. The studies were conducted in accordance with the local legislation and institutional requirements. The participants provided informed consent to participate in this study.

### Plasmids and Cell Lines

Codon-optimized, full-length open reading frames of SARS-CoV-2 spike genes (Table S1 and Fig. S1) were synthesized by GenScript (Piscataway, NJ, USA) and inserted into pVRC8400 (Vaccine Research Center, National Institutes of Health, Bethesda, MD). The HIV gag/pol packaging (pCMVΔR8.2) and firefly luciferase encoding transfer vector (pHR’CMV-Luc) plasmids [22, 23] were obtained from the Vaccine Research Center. 293T (ATCC, Manassas, VA; Cat no: CRL-11268), and 293T-ACE2-TMPRSS2 cells stably expressing human angiotensin-converting enzyme 2 (ACE2) and transmembrane serine protease 2 (TMPRSS2) (BEI Resources, Manassas, VA; Cat no: NR-55293) [24] were maintained at 37°C in Dulbecco’s modified eagle medium (DMEM) supplemented with high glucose, L-glutamine, minimal essential media (MEM) non-essential amino acids, penicillin/streptomycin, HEPES, and 10% fetal bovine serum (FBS).

### Cohort exposure groups

Participant characteristics and serum sample details are provided in Tables S2 and S3. Sera were collected from 25 healthy individuals at median 26 days (one month) and median 201 days (six months) after receiving the monovalent mRNA XBB.1.5 COVID-19 vaccine. All participants had previously received 3-4 doses of ancestral strain mRNA vaccine (WT) prior to receiving an XBB.1.5 vaccination during the 2023-2024 respiratory season. Participants were stratified into five groups based on COVID-19 vaccination and SARS-CoV-2 infection history (Table 1). The **WTs-XBB** group (n=2) included participants who received 3-4 WT vaccines followed by one XBB.1.5 vaccine, with no bivalent vaccination or SARS-CoV-2 infection. The **WTs-Bi-XBB** group (n=8) included participants who received 3-4 WT vaccines, one bivalent (ancestral/BA.4/5) (Bi) vaccine, and one XBB.1.5 vaccine, with no post-vaccination infection (PVI); one participant had a pre-vaccination D614G infection. The **WTs-BA.1PVI-Bi-XBB** group (n=6) included participants who received 3-4 WT vaccines, experienced a PVI during the BA.1 wave, and subsequently received one Bi vaccine and one XBB.1.5 vaccine. The **WTs-BA.4/5PVI-Bi-XBB** group (n=5) included participants who experienced a PVI during the BA.4/BA.5 wave prior to receipt of a Bi vaccine and subsequently received one or two Bi vaccines and one XBB.1.5 vaccine; vaccination histories varied (3 WT + 1 Bi, n=3; 4 WT + 2 Bi, n=1; 5 WT + 1 Bi, n=1). The **WTs-Bi-XBBPVI-XBB** group (n=4) included participants who received 3-4 WT vaccines and one Bi vaccine, experienced a PVI during the XBB wave, and subsequently received one XBB.1.5 vaccine. No participants had PCR-confirmed SARS-CoV-2 infection between XBB.1.5 vaccination and serum collection.

**Table 1.** XBB.1.5 booster antigenic exposure groups.

|  | Ancestral<br>vaccinations<br>(WT) | Bivalent<br>vaccinations<br>(ancestral/BA.4/5)<br>(Bi) | Post-vaccination<br>infection<br>(PVI)* | XBB.1.5<br>vaccination<br>(XBB) |
| --- | --- | --- | --- | --- |
| WTs-XBB, *n* = 2 | + | - | - | + |
| WTs-Bi-XBB, *n* = 8 | + | + | - | + |
| WTs-BA.1PVI-Bi-XBB, *n* = 6 | + | + | + BA.1 | + |
| WTs-BA.4/5PVI-Bi-XBB, *n* = 5 | + | + | + BA.4/5 | + |
| WTs-Bi-XBBPVI-XBB, *n* = 4 | + | + | + XBB | + |

\* Variant assigned based on the time of infection.
**WTs-XBB:** one participant with 3 WTs and 1 XBB; one participant with 4 WTs and 1 XBB.
**WTs-Bi-XBB:** four participants with 3 WTs, 1 Bi and 1 XBB; three participants with 4 WTs, 1 Bi and 1 XBB; one participant with 1 pre-vaccination D614G infection, 3 WTs, 1 Bi and 1 XBB.
**WTs-BA.1PVI-Bi-XBB:** four participants with 3 WTs, 1 BA.1 PVI, 1 Bi and 1 XBB; two participants with 4 WTs, 1 BA.1 PVI, 1 Bi and 1 XBB.
**WTs-BA.4/5PVI-Bi-XBB:** three participants with 3 WTs, 1 BA.4/5 PVI, 1 Bi and 1 XBB; one participant with 4 WTs, 1 BA.4/5 PVI, 2 Bi and 1 XBB; one participant with 5 WTs, 1 BA.4/5 PVI, 1 Bi and 1 XBB.
**WTs-Bi-XBBPVI-XBB:** three participants with 3 WTs, 1 Bi, 1 XBB PVI and 1 XBB; one participant with 4 WTs, 1 Bi, 1 XBB PVI and 1 XBB.

### Pseudovirus Neutralization Assay

Lentiviral pseudoviruses bearing SARS-CoV-2 spike proteins (Table S1) were generated as previously described [24]. 293T cells were co-transfected with 5 μg of pCMVΔR8.2, 5 μg of pHR’CMVLuc, and 0.5 μg of pVRC8400 encoding a codon-optimized spike gene. Supernatants were harvested ∼48 hours post transfection, filtered through a 0.45-μm low protein binding filter, and stored at −80°C. Neutralization assays were performed using 293T-ACE2-TMPRSS2 cells. Pseudoviruses with titers of ∼1 × 10^6^ relative luminescence units per milliliter (RLU/mL) were incubated with serially diluted sera for 2 hours at 37°C, then add to 96-well plates pre-seeded one day earlier with 3.0 × 10^4^ cells/well. After 48 hours infectivity was quantified by luciferase assay (Promega). Neutralization titer (ID_50_) was defined as the reciprocal serum dilution cause 50% RLU reduction relative to controls, calculated by nonlinear regression (GraphPad Prism Software, La Jolla, California). The geometric mean titer (GMT) from at least two independent experiments each with intra-assay duplicates was reported as the final titer.

### Statistical Analysis

Wilcoxon test for paired 2-group comparisons, Dunn’s multiple comparisons for multiple groups and GMTs with geometric SD were performed using GraphPad Prism. *P* values < 0.05 were considered significant. All neutralization titers were log2 transformed for analyses.

## RESULTS

### An additional mRNA XBB.1.5 vaccination after multiple prior COVID-19 vaccinations elicits broad neutralizing antibody responses across variants but reduced neutralization against newer variants

To investigate how prior antigenic exposure history shapes responses to new antigens, we analyzed sera collected one month after mRNA XBB.1.5 vaccination from individuals with diverse SARS-CoV-2 exposure histories. Participants were grouped into five antigenic exposure groups (Table 1): (1) WT vaccinations (WTs-XBB); (2) WT plus Bi vaccinations (WTs-Bi-XBB); (3) WT vaccinations, BA.1 PVI and Bi vaccination (WTs-BA.1PVI-Bi-XBB); (4) WT vaccinations, BA.4/ 5 PVI, and Bi vaccination (WTs-BA.4/5PVI-Bi-XBB); and (5) WT vaccinations, Bi vaccination, and XBB PVI (WTs-Bi-XBBPVI-XBB).

Neutralization titers were measured against pseudoviruses bearing spikes from pre-Omicron variants (D614G, B.1.617.2, and B.1.351), early Omicron variants (BA.1, BA.4/5, and XBB.1.5), late Omicron JN.1 variants (JN.1, KP.3, KP.3.1.1, XEC, LF.7, XFG, XFC, LP.8.1.1, and NB.1.8.1) that circulated after XBB.1.5, and divergent BA.3.2 variants (BA.3.2.1 and BA.3.2.2). Overall, one month post vaccination, sera exhibited high neutralization titers against pre-Omicron and early Omicron variants that significantly exceeded those against XBB.1.5 (Fig. 1A and Table S4). For example, titers against D614G (GMT = 14887) were over 8-fold greater than against XBB.1.5 (GMT = 1693), indicating strong back boosting to previously encountered antigens. Conversely, the XBB.1.5 vaccination elicited lower but detectable neutralization titers against the JN.1 lineage variants, with further reductions against variants containing additional escape mutations (KP.3, KP.3.1.1, XEC, LF.7, LP.8.1.1, NB.1.8.1, XFC and XFG). Interestingly, neutralization titers against JN.1 and the divergent BA.3.2 variants were similar.

**Figure 1.**
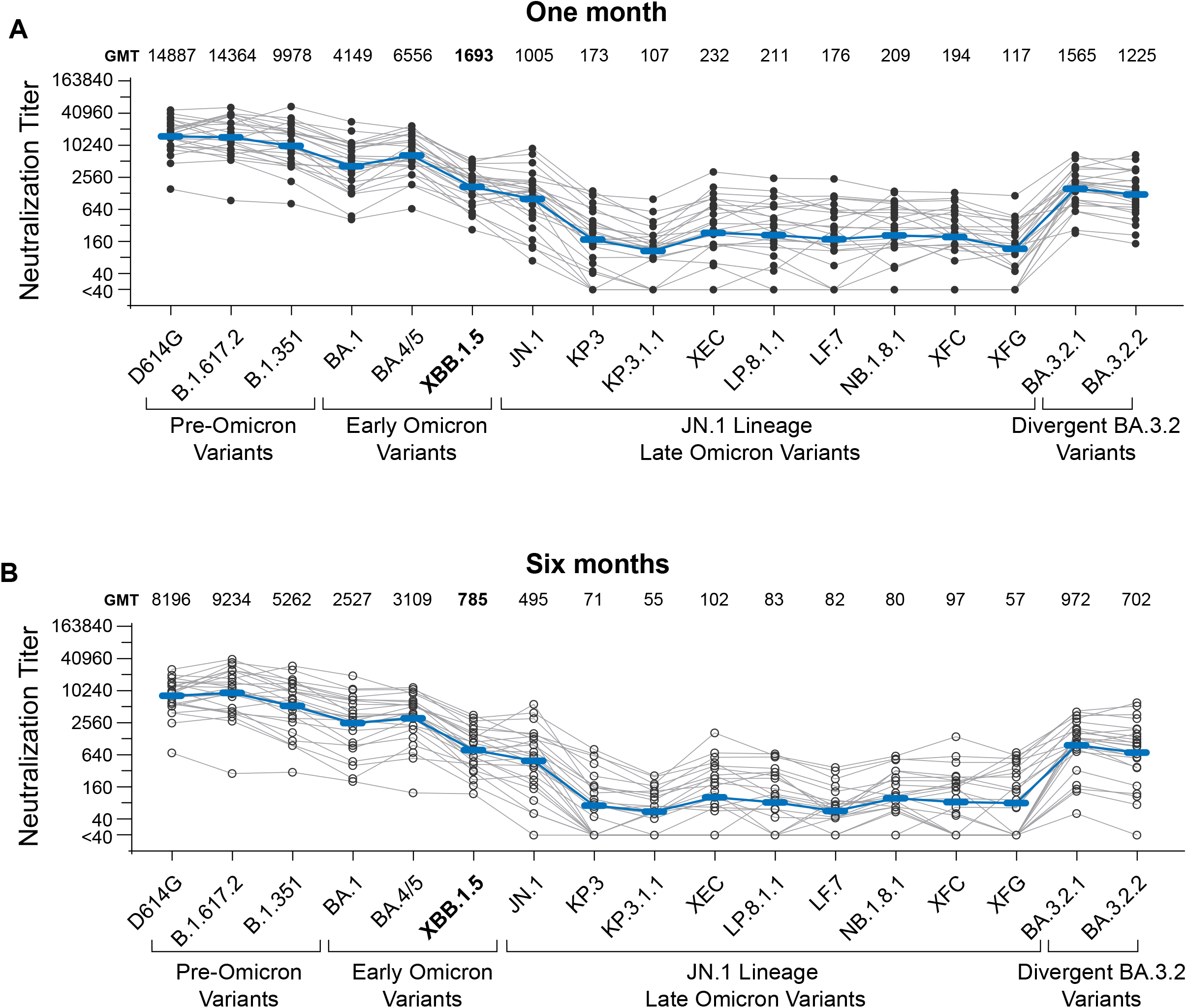
Neutralizing antibody responses following XBB.1.5 vaccination. **(A)** Serum neutralization titers against SARS-CoV-2 variants at one month post-XBB.1.5 vaccination across all exposure groups. **(B)** Serum neutralization titers against SARS-CoV-2 variants at six months post-XBB.1.5 vaccination across all exposure groups. Neutralization titers for each study participant against the indicated variants are plotted as a connected line. Blue lines connect geometric mean titers (GMT) for each variant. GMTs against individual variants are shown. Neutralization was measured using lentiviral-based pseudovirus neutralization assays. Serum samples with ID_50_ values below the limit of detection at 1:40 dilution were assigned an ID_50_ value of 20.

### Immunity wanes regardless of prior antigenic exposures, and titers are low against emerging variants

To investigate the long-term effects of repeated antigenic exposures on neutralizing antibody durability, we analyzed sera collected six months after XBB.1.5 vaccination from the same participants. At this time point, geometric mean titers (GMTs) of neutralization declined markedly across all variants (Figs. 1B and 2A, Table S5), with an average decrease of about 50%. Despite this universal decline, neutralization patterns were generally similar across all exposure groups (Fig. 2B), indicating that these differences in prior early Omicron antigenic exposures had limited influence on antibody durability following XBB.1.5 vaccination.

**Figure 2.**
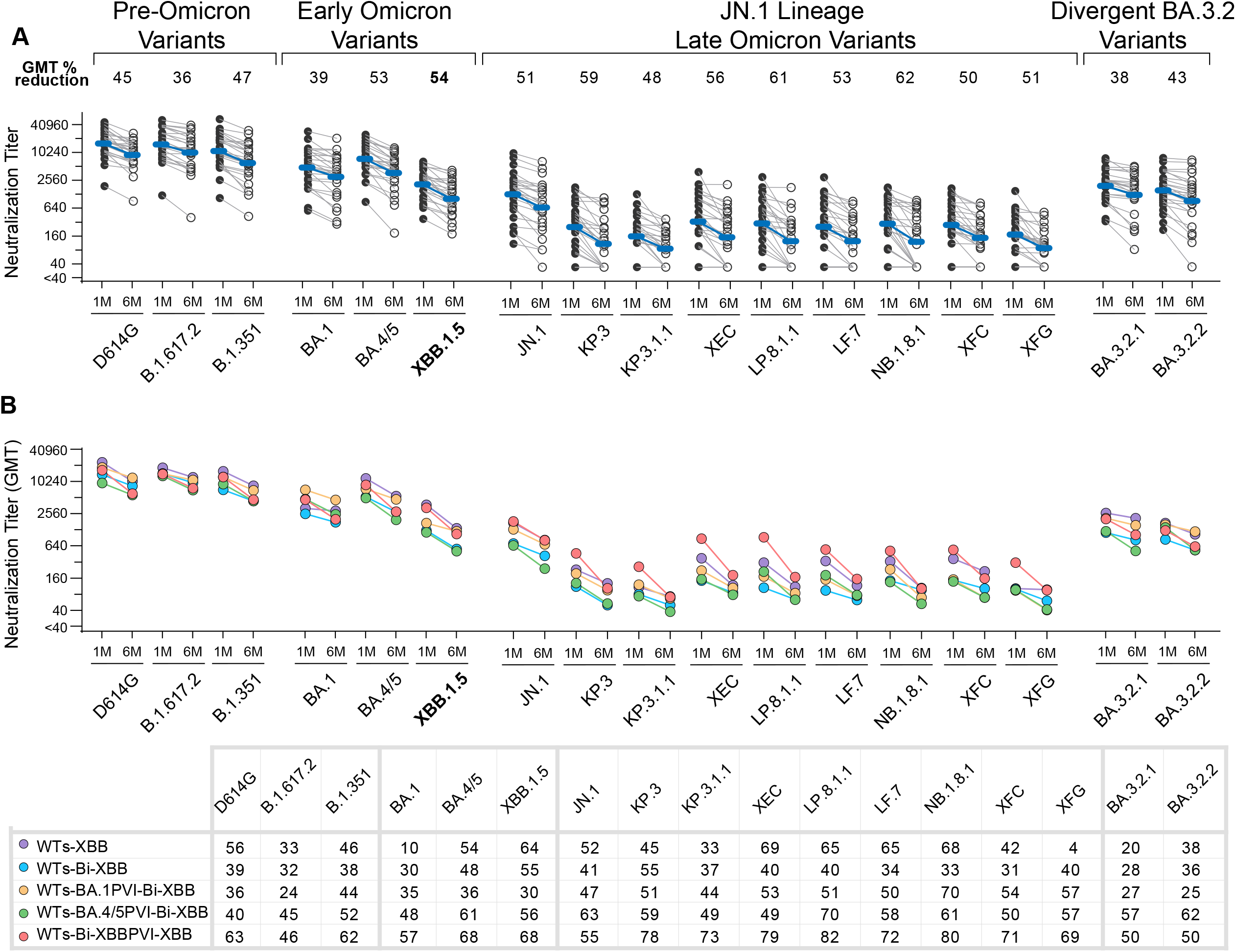
Neutralizing antibody durability across all exposure groups following XBB.1.5 vaccination. **(A)** Serum neutralization titers against SARS-CoV-2 variants at one month (1M) or six months (6M) post-XBB.1.5 vaccination across all exposure groups. Neutralization titers for each study participant against the indicated variants are plotted as a connected line. Blue lines connect geometric mean titers (GMT) at one and six months for each variant. GMT percentage reductions for each variant are indicated. **(B)** Neutralization GMT durability trends across exposure groups, showing neutralization reduction patterns against each tested variant at one month and six months post-vaccination. The table summarizes the GMT percentage reduction for each exposure group (sample details in Table1) (rows) across the panel of variants (columns). Neutralization was measured using lentiviral-based pseudovirus neutralization assays. Serum samples with ID_50_ values below the limit of detection at 1:40 dilution were assigned an ID_50_ value of 20.

**Figure 3.**
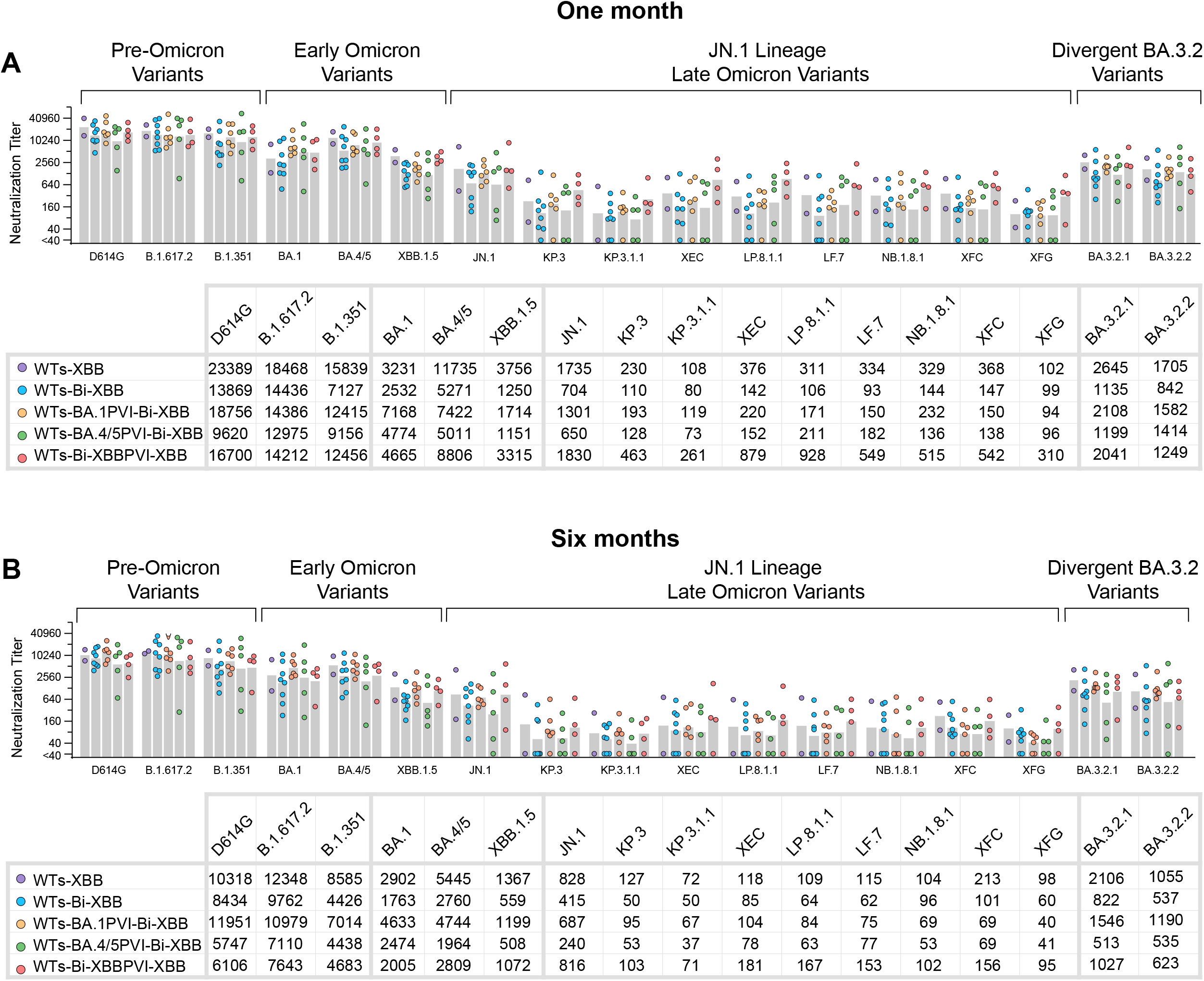
Neutralizing antibody responses in subjects with different prior exposure histories following XBB.1.5 vaccination. (**A**) Serum neutralization titers against SARS-CoV-2 variants at one month post-XBB.1.5 vaccination across different exposure groups. **(B)** Serum neutralization titers against SARS-CoV-2 variants at six months post-XBB.1.5 vaccination across different exposure groups. The tables below each graph summarize the GMTs for each exposure group (sample details in Table1) (rows) across the panel of variants (columns). Neutralization was measured using lentiviral-based pseudovirus neutralization assays. Serum samples with ID_50_ values below the limit of detection at 1:40 dilution were assigned an ID_50_ value of 20.

GMTs against the Delta variant (B.1.617.2) exhibited the smallest reduction (36%), while NB.1.8.1 (late Omicron variant) exhibited the largest (62%). Notably, GMTs targeting the vaccine strain, XBB.1.5, fell by 54%, consistent with the overall 50% average decrease across all variants.

Within an exposure group, the percent reduction in neutralizing antibodies was generally similar, (Fig. 2B and S2), though small sample sizes, particularly in the WTs-XBB group, limited definitive comparisons. The largest reductions (ranging from 46-82%) were observed in the WTs-Bi-XBBPVI-XBB group. Declines in the WTs-XBB, WTs-Bi-XBB, WTs-BA.1PVI-Bi-XBB, and WTs-BA.4/5PVI-Bi-XBB were less pronounced.

Overall, neutralization patterns against the variants were broadly consistent across exposure groups at both timepoints (Figs. 2B and 3). The WTs-Bi-XBB PVI-XBB group showed higher GMTs against JN.1 variants than groups without the extra XBB exposure, suggesting that a second XBB exposure broadened immune responses to the newer variants. Titers against many variants tended to be lower in the WTs-Bi-XBB, WTs-BA.1PVI-Bi-XBB, and WTs-BA.4/5PVI-Bi-XBB groups.

The absolute GMTs, which are important for assessing potentially protective antibody levels, differed substantially by variant across exposure groups. Despite the overall decline, absolute GMTs against pre-Omicron variants remained high at six months, with values of 8196 for D614G, 9234 for B.1.617.2, and 5262 for B.1.351 (Fig. 1B). In contrast, GMTs for early Omicron variants and BA.3.2 variants were lower at both timepoints in most groups, with a GMT for XBB.1.5 of 785 at six months post-vaccination. The late Omicron JN.1 sublineage variants, particularly those with enhanced immune evasion like KP.3, KP.3.1.1, XEC, LP.8.1.1, LF.7, NB.1.8.1, XFC, and XFG, had the lowest titers across all exposure groups, with GMTs falling below 110 at six months (Fig. 1B).

In summary, the XBB.1.5 vaccination elicited broad neutralizing responses in previously immunized participants, but titers waned substantially within six months, regardless of prior exposure history. Low neutralization titers against recently evolved variants at six months underscore the continuing challenge of maintaining protective immunity amid ongoing SARS-CoV-2 evolution.

### Neutralizing antibody responses to JN.1 sublineage variants after an mRNA XBB.1.5 in previously immunized participants are modestly influenced by N-glycan modifications in the spike N-terminal domain (NTD)

Because neutralization titers against KP.3, KP.3.1.1, XEC, LF.7, XFG, XFC, LP.8.1.1, and NB.1.8.1 variants were consistently lower than those against JN.1 across all exposure groups at both one- and six-month time points (Fig. 2), we investigated whether novel N-glycan modifications in the N-terminal domain (NTD) of spike proteins contribute to enhanced immune evasion in these variants.

Compared with KP.3 spike, KP.3.1.1 spike contains a serine deletion at residue 31 (DelS31) (Fig. S1), creating a new N-linked glycosylation site at position N30, which may function as a glycan shield that facilitates antibody escape [25]. In our cohort, neutralization titers against KP3.1.1 trended lower than those against KP.3 at both time points (Fig. 4A). Nearly all samples showed reduced neutralization titers against KP.3.1.1 compared with KP.3, though a few samples showed the opposite pattern. These results suggest that XBB.1.5 vaccination-induced neutralizing antibodies partially recognize epitopes influenced by S31 within the NTD.

**Figure 4.**
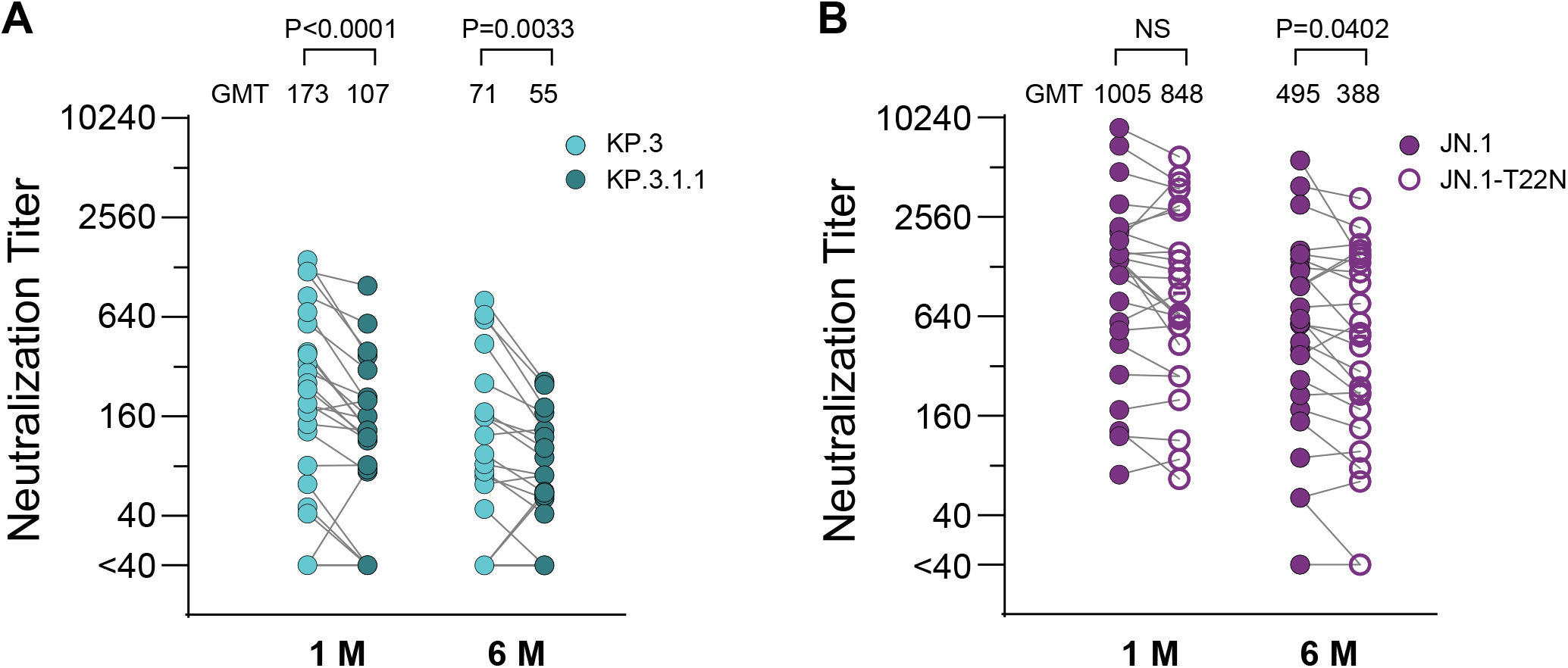
Impact of N-glycan modifications in spike N-terminal domain (NTD) on serum neutralization titers. Neutralization titer comparison between KP.3 and KP.3.1.1 variants in sera at one month and six months post-XBB.1.5 vaccination across overall exposure groups. **(B)** Neutralization titer comparison between JN.1 and JN.1 with T22N spike mutation in sera at one month and six months post-XBB.1.5 vaccination across overall exposure groups. Statistical comparisons between variants were analyzed using Wilcoxon test. *P* values <0.05 were considered statistically significant. NS, not significant. Neutralization was measured using lentiviral-based pseudovirus neutralization assays. Serum samples with ID_50_ values below the limit of detection at 1:40 dilution were assigned an ID_50_ value of 20.

The T22N mutation in some JN.1 sublineage variants (XEC, LF.7, XFG, XFC and NB.1.8.1) (Fig. S1) introduces an N-linked glycosylation sequon at position 22, resulting in glycan addition at this site. This glycan extends outward and partially overlaps a critical antibody epitope within the NTD [25]. To assess its effect, we engineered a T22N JN.1 spike variant. Neutralization titers against JN.1-T22N were generally lower than those against JN.1 at both time points (Fig. 4B). Over half the samples across exposure groups showed reduced titers against JN.1-T22N, while the remainder showed minimal or increased neutralization.

Together, these results suggest that N-glycan modifications within the NTD, such as DelS31 and T22N, contribute to the immune evasion of these newer variants. These findings indicate that XBB.1.5 vaccination-induced neutralizing antibodies partially recognize NTD epitopes, although we acknowledged our pseudovirus assay is more sensitive to antibodies targeting the receptor-binding site than those directed to the NTD.

## DISCUSSION

The rapid evolution of SARS-CoV-2, combined with heterogenous and waning population immunity, sustains COVID-19 as an ongoing public health concern. Understanding of how repeated antigenic exposures influence immunity to variants is needed to inform future vaccination strategies. This study evaluated neutralizing antibodies in a longitudinal cohort with well-documented COVID-19 vaccination and SARS-CoV-2 infection histories to assess whether repeated and varied antigenic exposures enhance the breadth and durability of humoral immunity. Although small group sizes limited statistical comparisons, our findings provide insights into how cumulative antigenic exposures influence variant immunity and vaccine responses, helping address key knowledge gaps given the rarity of large cohorts with well-defined exposure histories.

Despite the recognized advantages of hybrid immunity in enhancing antibody durability and neutralization breadth [17-19], we found that neither prior Omicron breakthrough infection nor prior bivalent vaccination substantially altered the magnitude or durability of neutralization responses after XBB.1.5 vaccination. This finding suggests that the benefits of hybrid immunity may become saturated after multiple exposures. Although additional vaccine doses reliably increased antibody titers, titers waned, and the ongoing emergence of immune-evasive variants supports continued consideration of updated vaccine formulations for at risk individuals.

Another key finding, consistent with previous reports, was strong evidence of immune imprinting (also called back boosting) [26, 27]. The XBB.1.5 vaccination elicited higher neutralizing antibody titers against previously encountered antigens (ancestral/D614G and early Omicron variants) than against vaccine-matched XBB.1.5. Such immune imprinting, often referred to as Original Antigenic Sin [28], reflects the dominance of memory responses from earlier exposures. This phenomenon was also observed in a pre-Omicron study where a Beta variant vaccine in individuals previously primed with the ancestral vaccine elicited higher neutralization titers against the ancestral variant than against the Beta variant [29].

Additional reports similarly described limited response to novel epitopes in BA.5 following bivalent vaccinations [30, 31], suggesting the dominance of immune memory focused on shared epitopes in previously seen antigens rather than *de novo* responses to novel Omicron epitopes. A recent report further showed that individuals without prior bivalent vaccination exhibited greater boosting of neutralizing antibodies against XBB.1.5 and JN.1 after XBB.1.5 vaccination than those previously given bivalent vaccines, suggesting a dampening effect of prior bivalent exposure [32]. However, it remains unclear whether memory responses actively inhibit responses to novel epitopes. Memory responses could hinder responses to novel epitopes through mechanisms like accelerated antigen clearance or epitope masking, or they may simply dominate through superior memory recall and higher-affinity responses without directly suppressing primary responses. Nonetheless, a booster vaccination increases neutralizing antibody titers overall, irrespective of whether the boosted antibodies target conserved or novel epitopes. However, eliciting responses to a new variant’s novel epitopes would have the advantage of priming a rapid memory response for any subsequent exposure to that variant.

A related observation was that participants with two XBB exposures (XBB infection and vaccination) had the highest titers against XBB.1.5, suggesting a second exposure broadened responses to novel XBB.1.5 epitopes [33]. Participants with a long interval between their ancestral and XBB.1.5 vaccinations also had relatively high titers, consistent with findings that extended intervals between antigenic exposures enhance responses to novel antigens [34-36]. These observations underscore the importance of updating vaccine to enhance responses to new variants, while preserving the benefits of back boosting.

At six months post XBB.1.5 vaccination, GMTs declined by 36–62% across all variants and exposure groups, reflecting the natural contraction of humoral immunity following acute antigenic exposure that is dominated by short-lived plasma cells. The WTs-Bi-XBBPVI-XBB group exhibited the largest declines (46–82%), particularly against JN.1 variants, suggesting that homologous boosting, while generating strong peak responses, may not establish long-lived memory B cells capable of sustained high-level neutralizing antibodies.

Imprinting remained prominent at six months. While titers against the ancestral strain (D614G) remained relatively high, the lowest titers at both one and six months were consistently against the JN.1 sublineage variants, confirming their enhanced immune evasions. A recent study reported waning patterns after XBB.1.5 boosting, but breadth against some of the newer variants was not assessed [26]. The rapid decline of cross-neutralization against highly evasive JN.1 variants highlights the difficulty of sustaining broad protection.

Consistent reduction in neutralization across JN.1 lineage variants harboring mutations that alter glycosylation in the NTD led us to investigate these glycan modifications as potential immune evasion mechanisms. The DelS31 mutation in KP.3.1.1, which has been observed in other sarbecoviruses [37], introduces a glycan at N30 [25], while the T22N mutation creates a new glycosylation site [38]. Sera showed slightly lower neutralization against these NTD-modified variants, suggesting that XBB.1.5 boosting in this cohort primarily induced neutralizing antibodies that target epitopes outside the NTD.

Evolution from BA.2.86 to JN.1 and its JN.1 sublineages reflects progressive acquisition of recurrent mutations, including R346T, L455S, F456L, and Q493E that collectively enhance viral fitness and antibody escape, with L455S playing a major role [33]. The recurrent ancestral residues, such as the N30 glycan and Q493E mutation, also found in other sarbecoviruses, suggests evolutionary optimization balancing immune escape with viral fitness [37].

Interestingly, XBB.1.5 vaccination elicited neutralization against BA.3.2 at levels comparable to XBB.1.5 and JN.1, with similar durability to early Omicron variants. Although BA.3.2 currently appears to lack replication fitness relative to some other variants [39], its 44 unique spike mutations compared with LP.8.1.1 that is included in recently updated vaccines (Fig. S1B), raise concern for potential antigenic shift. However, it still shares approximately 19 mutations with XBB.1.5 and JN.1 sublineages (Fig. S1B). Recent studies show that KP.2-formulated COVID-19 vaccination elicits comparable neutralization of BA.3.2 and JN.1 [40], while others report limited BA.3.2 neutralization after BA.5 and JN.1 exposure [39]. The higher titers observed in our cohort may reflect the participants’ extensive and distinct antigenic exposure histories, assay differences, or both.

Overall, the rapid decline in cross-neutralizing antibodies and persistent immune imprinting reinforce the challenge of maintaining durable immunity with current monovalent mRNA vaccines and underscore the potential benefit of updating vaccine formulation for eligible persons. The emergence of variants with highly balanced immune evasive and fitness profiles emphasizes the need for continuous, antigenic surveillance and real-time data to guide these updates. The marked reduction in titers against highly evasive variants supports considerations for timely boosting to maintain protective antibody levels, especially for high-risk individuals. Long-term success will likely require new vaccination strategies, such as novel antigenic designs or alternative delivery platforms, to improve neutralization breadth and durability, as well as enhance mucosal immunity.

This study has several limitations, including small subgroup sizes, uncontrolled timing intervals between antigenic exposures, lack of pre-XBB.1.5 vaccination sera, and assessment limited to humoral responses. Our pseudovirus assay is less sensitive to NTD-focused antibodies than those targeting the receptor binding site. Finally, the six months follow-up period precludes conclusions about long-term memory B-cell durability.

In summary, in individuals with extensive prior COVID-19 antigenic exposures, monovalent mRNA XBB.1.5 vaccination generated broadly neutralizing antibodies heavily influenced by immunological imprinting, with substantial waning within six months. Lower peak titers against the most antigenically divergent JN.1 sublineages resulted in very low neutralizing titers at six months, suggesting limited residual antibody protection. Additional antigenic exposures added minimal durability or breadth against immune-evasive variants. The marked decline in neutralization against newer variants over typical booster intervals underscores the need for ongoing antigenic surveillance and potentially strategic refinement of booster timing and vaccine composition to maintain protection against evolving variants, especially for populations at highest risk.

## Figure legends

**Figure S1. SARS-CoV-2 spike mutations across XBB.1.5, BA.3.2, and JN.1 sublineage variants. (A)** Amino acid mutations and deletions (Del) in spike proteins of JN.1 sublineage variants relative to JN.1 reference sequence. Blue boxes indicate amino acid substitutions relative to JN.1. **(B)** Venn diagram of shared and unique mutations between XBB.1.5, BA.3.2.1, BA.3.2.2, JN.1, and LP.8.1.1. Red residues highlight amino acid positions that are mutated in multiple variants.

**Figure S2. Neutralizing antibody durability comparison across all exposure groups following XBB.1.5 vaccination**. Comparative neutralization durability trends across all exposure groups (sample details in Table1), showing neutralization reduction patterns against each tested variant at one month and six months post-vaccination. Individual participant data are plotted as dots. Reduction % GMT ratios (one month vs. six months) against each tested variant in each exposure group are shown. Neutralization was measured using lentiviral-based pseudovirus neutralization assays. Serum samples with ID_50_ values below the limit of detection at 1:40 dilution were assigned an ID_50_ value of 20.

## Acknowledgments

The investigators gratefully acknowledge all research participants for their many contributions to the PASS study.

## Author disclaimer

The views expressed in this presentation are the sole responsibility of the presenter and do not necessarily reflect the views, opinions, or policies of the Uniformed Services University of the Health Sciences, the Department of Defense, the Department of the Navy, Army, Air Force, Defense Health Agency, US Food and Drug Administration, the United States Government, or the Henry M. Jackson Foundation for the Advancement of Military Medicine, Inc. (HJF). Several of the authors are U.S. Government employees. This work was prepared as part of their official duties. Title 17 US.C. § 105 provides that “Copyright protection under this title is not available for any work of the United States Government”. Title 17 US.C. § 101 defines a U.S. Government work as a work prepared by a military service member or employee of the U.S. Government as part of that person’s official duties. Mention of trade names, commercial products, or organizations does not imply endorsement by the U.S. Government. The study protocol was approved by the USU Institutional Review Board in compliance with all applicable federal regulations governing the protection of human subjects.

## REFERENCES

1. Jian F, Wang J, Yisimayi A, et al. Evolving antibody response to SARS-CoV-2 antigenic shift from XBB to JN.1. Nature 2025; 637:921–9.

2. Planas D, Staropoli I, Michel V, et al. Distinct evolution of SARS-CoV-2 Omicron XBB and BA.2.86/JN.1 lineages combining increased fitness and antibody evasion. Nat Commun 2024; 15:2254.

3. Wang Q, Guo Y, Iketani S, et al. Antibody evasion by SARS-CoV-2 Omicron subvariants BA.2.12.1, BA.4 and BA.5. Nature 2022; 608:603–8.

4. Wu Q, Wu H, Hu Y, et al. Immune evasion of Omicron variants JN.1, KP.2, and KP.3 to the polyclonal and monoclonal antibodies from COVID-19 convalescents and vaccine recipients. Antiviral Res 2025; 235:106092.

5. World Health Organization. Types of data requested to inform December 2025 COVID-19 vaccine antigen composition deliberations. Available at: https://www.who.int/news/item/29-09-2025-types-of-data-requested-to-inform-december-2025-covid-19-vaccine-antigen-composition-deliberations. Accessed 30 September 2025.

6. Khoury DS, Cromer D, Reynaldi A, et al. Neutralizing antibody levels are highly predictive of immune protection from symptomatic SARS-CoV-2 infection. Nat Med 2021; 27:1205–11.

7. Zheng Y, Pan J, Jin M, et al. Efficacy of the neutralizing antibodies after the booster dose on SARS-CoV-2 Omicron variant and a two-year longitudinal antibody study on Wild Type convalescents. Int Immunopharmacol 2023; 119:110151.

8. Ou S, Huang Z, Lan M, et al. The duration and breadth of antibody responses to 3-dose of inactivated COVID-19 vaccinations in healthy blood donors: An observational study. Front Immunol 2022; 13:1027924.

9. Srivastava K, Carreno JM, Gleason C, et al. SARS-CoV-2-infection- and vaccine-induced antibody responses are long lasting with an initial waning phase followed by a stabilization phase. Immunity 2024; 57:587–99 e4.

10. Rosati M, Terpos E, Ntanasis-Stathopoulos I, et al. Sequential Analysis of Binding and Neutralizing Antibody in COVID-19 Convalescent Patients at 14 Months After SARS-CoV-2 Infection. Front Immunol 2021; 12:793953.

11. Steiner LA, Eisen HN. The relative affinity of antibodies synthesized in the secondary response. J Exp Med 1967; 126:1185–205.

12. Siskind GW, Benacerraf B. Cell selection by antigen in the immune response. Adv Immunol 1969; 10:1–50.

13. Gitlin AD, Shulman Z, Nussenzweig MC. Clonal selection in the germinal centre by regulated proliferation and hypermutation. Nature 2014; 509:637–40.

14. Collier DA, Ferreira I, Kotagiri P, et al. Age-related immune response heterogeneity to SARS-CoV-2 vaccine BNT162b2. Nature 2021; 596:417–22.

15. Goel RR, Apostolidis SA, Painter MM, et al. Distinct antibody and memory B cell responses in SARS-CoV-2 naive and recovered individuals following mRNA vaccination. Sci Immunol 2021; 6.

16. Muecksch F, Weisblum Y, Barnes CO, et al. Affinity maturation of SARS-CoV-2 neutralizing antibodies confers potency, breadth, and resilience to viral escape mutations. Immunity 2021; 54:1853–68 e7.

17. Voss WN, Mallory MA, Byrne PO, et al. Hybrid immunity to SARS-CoV-2 arises from serological recall of IgG antibodies distinctly imprinted by infection or vaccination. Cell Rep Med 2024; 5:101668.

18. Reynolds CJ, Pade C, Gibbons JM, et al. Prior SARS-CoV-2 infection rescues B and T cell responses to variants after first vaccine dose. Science 2021; 372:1418–23.

19. Manali M, Bissett LA, Amat JAR, et al. SARS-CoV-2 Evolution and Patient Immunological History Shape the Breadth and Potency of Antibody-Mediated Immunity. J Infect Dis 2022; 227:40–9.

20. Pusnik J, Monzon-Posadas WO, Zorn J, et al. SARS-CoV-2 humoral and cellular immunity following different combinations of vaccination and breakthrough infection. Nat Commun 2023; 14:572.

21. Jackson-Thompson BM, Goguet E, Laing ED, et al. Prospective Assessment of SARS-CoV-2 Seroconversion (PASS) study: an observational cohort study of SARS-CoV-2 infection and vaccination in healthcare workers. BMC Infect Dis 2021; 21:544.

22. Naldini L, Blomer U, Gallay P, et al. In vivo gene delivery and stable transduction of nondividing cells by a lentiviral vector. Science 1996; 272:263–7.

23. Zufferey R, Nagy D, Mandel RJ, Naldini L, Trono D. Multiply attenuated lentiviral vector achieves efficient gene delivery in vivo. Nat Biotechnol 1997; 15:871–5.

24. Neerukonda SN, Vassell R, Herrup R, et al. Establishment of a well-characterized SARS-CoV-2 lentiviral pseudovirus neutralization assay using 293T cells with stable expression of ACE2 and TMPRSS2. PLoS One 2021; 16:e0248348.

25. Li P, Faraone JN, Hsu CC, et al. Neutralization and spike stability of JN.1-derived LB.1, KP.2.3, KP.3, and KP.3.1.1 subvariants. mBio 2025; 16:e0046425.

26. Kumar S, Jain S, Wali B, et al. The XBB.1.5 COVID-19 vaccine elicits a durable antibody response to ancestral and XBB.1.5 SARS-CoV-2 spike proteins. Sci Transl Med 2025; 17:eadu8067.

27. Roltgen K, Nielsen SCA, Silva O, et al. Immune imprinting, breadth of variant recognition, and germinal center response in human SARS-CoV-2 infection and vaccination. Cell 2022; 185:1025–40 e14.

28. Francis T. On the Doctrine of Original Antigenic Sin. Proceedings of the American Philosophical Society 1960; 104:572–8.

29. Wheatley AK, Fox A, Tan HX, et al. Immune imprinting and SARS-CoV-2 vaccine design. Trends Immunol 2021; 42:956–9.

30. Collier AY, Miller J, Hachmann NP, et al. Immunogenicity of BA.5 Bivalent mRNA Vaccine Boosters. N Engl J Med 2023; 388:565–7.

31. Wang Q, Bowen A, Valdez R, et al. Antibody Response to Omicron BA.4-BA.5 Bivalent Booster. N Engl J Med 2023; 388:567–9.

32. Wrynla XH, Bates TA, Trank-Greene M, et al. Immune imprinting and vaccine interval determine antibody responses to monovalent XBB.1.5 COVID-19 vaccination. Commun Med (Lond) 2025; 5:182.

33. Li P, Faraone JN, Hsu CC, et al. Neutralization escape, infectivity, and membrane fusion of JN.1-derived SARS-CoV-2 SLip, FLiRT, and KP.2 variants. Cell Reports 2024; 43.

34. Bates TA, Leier HC, McBride SK, et al. An extended interval between vaccination and infection enhances hybrid immunity against SARS-CoV-2 variants. JCI Insight 2023; 8.

35. Payne RP, Longet S, Austin JA, et al. Immunogenicity of standard and extended dosing intervals of BNT162b2 mRNA vaccine. Cell 2021; 184:5699–714 e11.

36. Grunau B, Goldfarb DM, Asamoah-Boaheng M, et al. Immunogenicity of Extended mRNA SARS-CoV-2 Vaccine Dosing Intervals. JAMA 2022; 327:279–81.

37. Feng Z, Huang J, Baboo S, et al. Structural and Functional Insights into the Evolution of SARS-CoV-2 KP.3.1.1 Spike Protein. bioRxiv 2024.

38. Li P, Faraone JN, Hsu CC, et al. Role of glycosylation mutations at the N-terminal domain of SARS-CoV-2 XEC variant in immune evasion, cell-cell fusion, and spike stability. J Virol 2025; 99:e0024225.

39. Guo C, Yu Y, Liu J, et al. Antigenic and virological characteristics of SARS-CoV-2 variants BA.3.2, XFG, and NB.1.8.1. Lancet Infect Dis 2025; 25:e374–e7.

40. Abbad A, Lerman B, Ehrenhaus J, et al. Antibody responses to SARS-CoV-2 variants LP.8.1, LF.7.1, NB.1.8.1, XFG and BA.3.2 following KP.2 monovalent mRNA vaccination. medRxiv 2025.

