## Supplemental material for "Repeated SARS-CoV-2 Antigenic Exposures from Prior Vaccinations and Infections Demonstrate Limits of Antibody Durability and Breadth Against Newer Variants"

Figure S1

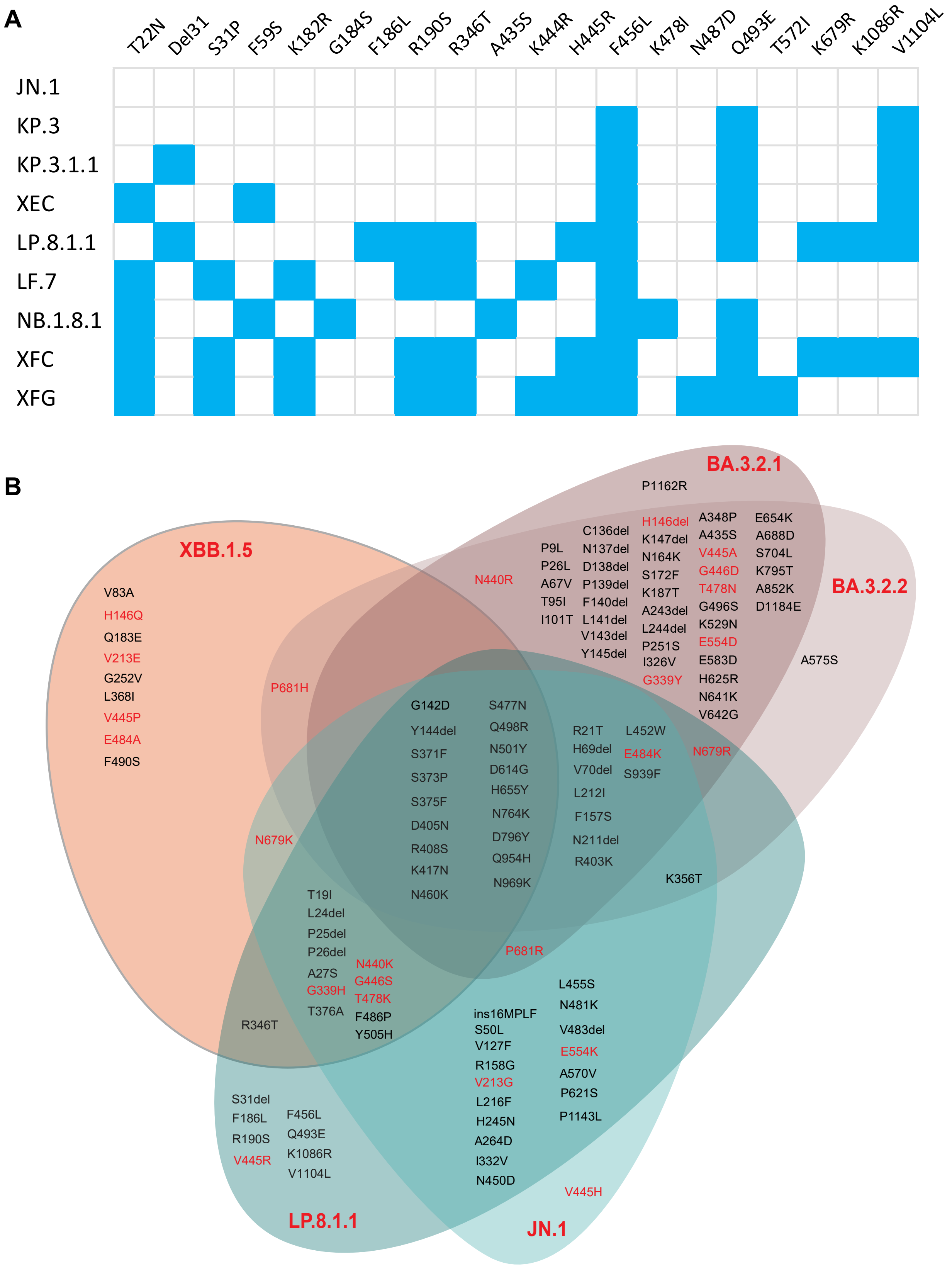
A
T22N
Del31
S31P
F59S
K182R
G184S
F186L
R190S
R346T
A435S
K444R
H445R
F456L
K478I
N487D
Q493E
T572I
K679R
K1086R
V1104L
JN.1
KP.3
KP.3.1.1
XEC
LP.8.1.1
LF.7
NB.1.8.1
XFC
XFG
B
BA.3.2.1
BA.3.2.2
XBB.1.5
LP.8.1.1
JN.1
P1162R
H146del
K147del
N164K
S172F
K187T
A243del
L244del
P251S
I326V
G339Y
A348P
A435S
V445A
G446D
T478N
G496S
K529N
E554D
E583D
H625R
N641K
V642G
E654K
A688D
S704L
K795T
A852K
D1184E
C136del
N137del
D138del
P139del
F140del
L141del
V143del
Y145del
P9L
P26L
A67V
T95I
I101T
N440R
A575S
V83A
H146Q
Q183E
V213E
G252V
L368I
V445P
E484A
F490S
P681H
G142D
Y144del
S371F
S373P
S375F
D405N
R408S
K417N
N460K
S477N
Q498R
N501Y
D614G
H655Y
N764K
D796Y
Q954H
N969K
R21T
H69del
V70del
L212I
F157S
N211del
R403K
L452W
E484K
S939F
N679R
N679K
K356T
T19I
L24del
P25del
P26del
A27S
G339H
T376A
N440K
G446S
T478K
F486P
Y505H
P681R
R346T
L455S
N481K
V483del
E554K
A570V
P621S
P1143L
ins16MPLF
S50L
V127F
R158G
V213G
L216F
H245N
A264D
I332V
N450D
S31del
F186L
R190S
V445R
F456L
Q493E
K1086R
V1104L
V445H

### Figure S2

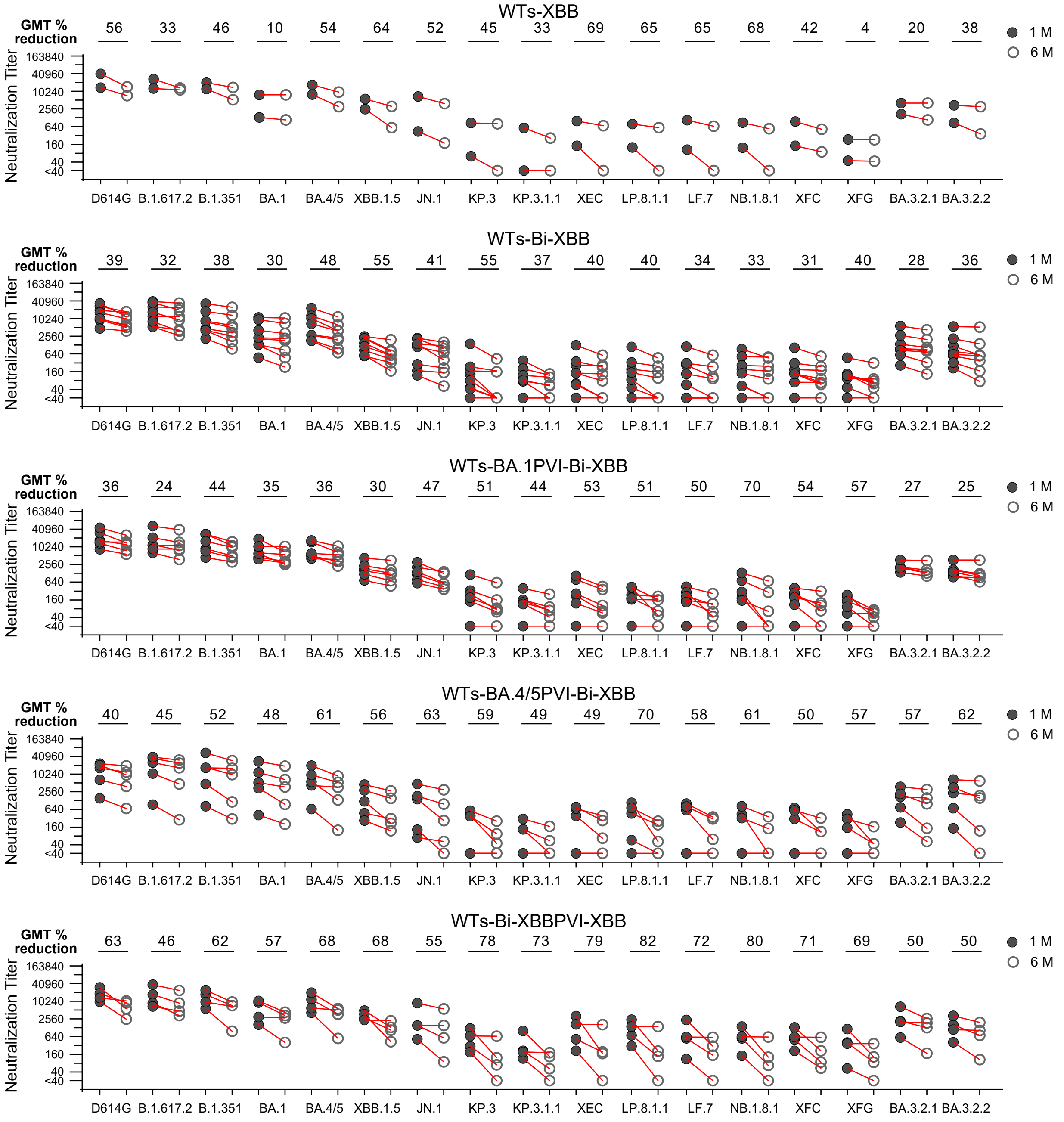
WTs-XBB
GMT % reduction
56 33 46 10 54 64 52 45 33 69 65 65 68 42 4 20 38
Neutralization Titer
163840 40960 10240 2560 640 160 40 <40
D614G B.1.617.2 B.1.351 BA.1 BA.4/5 XBB.1.5 JN.1 KP.3 KP.3.1.1 XEC LP.8.1.1 LF.7 NB.1.8.1 XFC XFG BA.3.2.1 BA.3.2.2
1 M
6 M
WTs-Bi-XBB
GMT % reduction
39 32 38 30 48 55 41 55 37 40 40 34 33 31 40 28 36
Neutralization Titer
163840 40960 10240 2560 640 160 40 <40
D614G B.1.617.2 B.1.351 BA.1 BA.4/5 XBB.1.5 JN.1 KP.3 KP.3.1.1 XEC LP.8.1.1 LF.7 NB.1.8.1 XFC XFG BA.3.2.1 BA.3.2.2
1 M
6 M
WTs-BA.1PVI-Bi-XBB
GMT % reduction
36 24 44 35 36 30 47 51 44 53 51 50 70 54 57 27 25
Neutralization Titer
163840 40960 10240 2560 640 160 40 <40
D614G B.1.617.2 B.1.351 BA.1 BA.4/5 XBB.1.5 JN.1 KP.3 KP.3.1.1 XEC LP.8.1.1 LF.7 NB.1.8.1 XFC XFG BA.3.2.1 BA.3.2.2
1 M
6 M
WTs-BA.4/5PVI-Bi-XBB
GMT % reduction
40 45 52 48 61 56 63 59 49 49 70 58 61 50 57 57 62
Neutralization Titer
163840 40960 10240 2560 640 160 40 <40
D614G B.1.617.2 B.1.351 BA.1 BA.4/5 XBB.1.5 JN.1 KP.3 KP.3.1.1 XEC LP.8.1.1 LF.7 NB.1.8.1 XFC XFG BA.3.2.1 BA.3.2.2
1 M
6 M
WTs-Bi-XBBPVI-XBB
GMT % reduction
63 46 62 57 68 68 55 78 73 79 82 72 80 71 69 50 50
Neutralization Titer
163840 40960 10240 2560 640 160 40 <40
D614G B.1.617.2 B.1.351 BA.1 BA.4/5 XBB.1.5 JN.1 KP.3 KP.3.1.1 XEC LP.8.1.1 LF.7 NB.1.8.1 XFC XFG BA.3.2.1 BA.3.2.2
1 M
6 M

**Table S1. SARS-CoV-2 spikes used for pseudoviruses.**

| Virus name | GenBank or GISAID # |
|---|---|
| D614G | EPI_ISL_5851484 |
| B.1.617.2 | MW934201 |
| B.1.351 | MW598419 |
| BA.1 | OQ361639 |
| BA.4/5 | EPI_ISL_12464782 |
| XBB.1.5 | EPI_ISL_15687648 |
| JN.1 | OR649559 |
| KP.3 | PP599813 |
| KP.3.1.1 | PP897603 |
| XEC | PQ147446 |
| LF.7 | EPI_ISL_19707256 |
| XFG | EPI_ISL_19724626 |
| XFC | EPI_ISL_19658767 |
| LP.8.1.1 | EPI_ISL_19808041 |
| NB.1.8.1 | EPI_ISL_19750627 |
| BA.3.2.1 | EPI_ISL_19771107 |
| BA.3.2.2 | EPI_ISL_19771105 |

**Table S2. Summary of antigenic exposures for each participant**

| Sample ID | COVID-19 vaccination fall 2023 | Number of vaccinations | COVID-19 infection date (month-year) | XBB.1.5 Booster date (month-year) | Days since XBB.1.5 booster vaccination (1 month) | Days since XBB.1.5 booster vaccination (6 months) |
|---|---|---|---|---|---|---|
| 1 | **Pfizer** | **5** | | Sep-25 | 35 | 197 |
| 2 | **Pfizer** | **4** | | Nov-25 | 11 | 223 |
| 3 | **Pfizer** | **6** | | Nov-25 | 30 | 198 |
| 4 | **Pfizer** | **5** | | Oct-25 | 22 | 227 |
| 5 | **Moderna** | **5** | | Nov-25 | 9 | 190 |
| 6 | **Pfizer** | **6** | | Oct-25 | 28 | 203 |
| 7 | **Pfizer** | **5** | Nov-25 | Nov-25 | 7 | 159 |
| 8 | **Pfizer** | **5** | | Oct-25 | 52 | 241 |
| 9 | **Pfizer** | **6** | | Oct-25 | 40 | 166 |
| 10 | **Pfizer** | **5** | | Oct-25 | 23 | 177 |
| 11 | **Pfizer** | **5** | Mar-25 | Nov-25 | 32 | 158 |
| 12 | **Pfizer** | **5** | Feb-25 | Oct-25 | 20 | 202 |
| 13 | **Pfizer** | **6** | Jan-25 | Sep-25 | 46 | 186 |
| 14 | **Pfizer** | **5** | Dec-25 | Oct-25 | 28 | 238 |
| 15 | **Pfizer** | **6** | Dec-25 | Oct-25 | 20 | 248 |
| 16 | **Pfizer** | **5** | Jan-25 | Oct-25 | 16 | 191 |
| 17 | **Moderna** | **7** | Apr-25 | Sep-25 | 56 | 202 |
| 18 | **Moderna** | **7** | May-25 | Oct-25 | 11 | 188 |
| 19 | **Pfizer** | **5** | May-25 | Oct-25 | 26 | 201 |
| 20 | **Pfizer** | **5** | May-25 | Oct-25 | 36 | 191 |
| 21 | **Moderna** | **5** | Jun-25 | Oct-25 | 27 | 215 |
| 22 | **Pfizer** | **5** | Oct-25 | Oct-25 | 21 | 204 |
| 23 | **Pfizer** | **5** | Nov-25 | Oct-25 | 28 | 203 |
| 24 | **Pfizer** | **5** | Jul-25 | Oct-25 | 16 | 162 |
| 25 | **Pfizer** | **6** | Dec-25 | Oct-25 | 21 | 202 |

### Table S3. Demographic information of the antigenic exposure groups

| | | Overall | WTs-XBB | WTs-Bi-XBB | WTs-BA.1PVI-Bi-XBB | WTs-BA.4/5PVI-Bi-XBB | WTs-Bi-XBBPVI-XBB |
|---|---|---|---|---|---|---|---|
| | | N=25 | N=2 | N=8 | N=6 | N=5 | N=4 |
| **Age group** | | | | | | | |
| | 18-44 | 9 (36.0%) | 1 (50.0%) | 2 (22.2%) | 2 (33.3%) | 2 (40.0%) | 2 (50.0%) |
| | 45-64 | 15 (60.0%) | 0 (0%) | 7 (77.8%) | 3 (50.0%) | 3 (60.0%) | 2 (50.0%) |
| | 65+ | 2 (8.0%) | 1 (50.0%) | 0 (0%) | 1 (16.7%) | 0 (0%) | 0 (0%) |
| | Median age (IQR) | 47 (39 - 57) | 54 (45 - 63) | 49 (45 - 55) | 46 (41 - 53) | 57 (43 - 62) | 45 (38 - 54) |
| **Gender** | | | | | | | |
| | Female | 17 (68.0%) | 2 (100%) | 6 (75.0%) | 4 (66.7%) | 2 (40.0%) | 3 (75.0%) |
| | Male | 8 (32.0%) | 0 (0%) | 2 (25.0%) | 2 (33.3%) | 3 (60.0%) | 1 (25.0%) |
| **Race** | | | | | | | |
| | White | 18 (72.0%) | 2 (100%) | 4 (50.0%) | 5 (83.3%) | 4 (80.0%) | 3 (75.0%) |
| | Black | 3 (12.0%) | 0 (0%) | 2 (25.0%) | 0 (0%) | 0 (0%) | 1 (25.0%) |
| | Asian | 4 (16.0%) | 0 (0%) | 2 (25.0%) | 1 (16.7%) | 1 (20.0%) | 0 (0%) |
| **Ethnicity** | | | | | | | |
| | Hispanic | 1 (4.0%) | 0 (0%) | 0 (0%) | 0 (0%) | 0 (0%) | 1 (25.0%) |
| | Non Hispanic | 24 (96.0%) | 2 (100%) | 8 (100%) | 6 (100%) | 5 (100%) | 3 (75.0%) |
| **Primary vaccine series** | | | | | | | |
| | Pfizer BNT162b2 | 25 (100%) | 2 (100%) | 8 (100%) | 6 (100%) | 5 (100%) | 4 (100%) |
| **First booster (after primary series)** | | | | | | | |
| | Pfizer BNT162b2 | 24 (96.0%) | 2 (100%) | 8 (100%) | 6 (100%) | 4 (80.0%) | 4 (100%) |
| | Moderna mRNA-1273 | 1 (4.0%) | 0 (0%) | 0 (0%) | 0 (0%) | 1 (20.0%) | 0 (0%) |
| **Second booster** | | | | | | | |
| | Pfizer BNT162b2 | 9 (36.0%) | 1 (50.0%) | 3 (37.5%) | 2 (33.3%) | 2 (40.0%) | 1 (25.0%) |
| | Pfizer bivalent (original and Omicron BA.4/BA.5) | 14 (56.0%) | 0 (0%) | 5 (62.5%) | 4 (66.7%) | 2 (40.0%) | 3 (75.0%) |
| | Moderna bivalent (original and Omicron BA.4/BA.5) | 1 (4.0%) | 0 (0%) | 0 (0%) | 0 (0%) | 1 (20.0%) | 0 (0%) |
| | Pfizer monovalent XBB.1.5 | 1 (4.0%) | 1 (50.0%) | 0 (0%) | 0 (0%) | 0 (0%) | 0 (0%) |
| **Third booster** | | | | | | | |
| | Pfizer BNT162b2 | 1 (4.0%) | 0 (0%) | 0 (0%) | 0 (0%) | 1 (20.0%) | 0 (0%) |
| | Pfizer bivalent (original and Omicron BA.4/BA.5) | 7 (28.0%) | 0 (0%) | 3 (37.5%) | 2 (33.3%) | 1 (20.0%) | 1 (25.0%) |
| | Pfizer monovalent XBB.1.5 | 15 (60.0%) | 1 (50.0%) | 5 (62.5%) | 4 (66.7%) | 2 (40.0%) | 3 (75.0%) |
| | Moderna monovalent XBB.1.5 | 1 (4.0%) | 0 (0%) | 0 (0%) | 0 (0%) | 1 (20.0%) | 0 (0%) |
| | Unboosted | 1 (4.0%) | 1 (50.0%) | 0 (0%) | 0 (0%) | 0 (0%) | 0 (0%) |
| **Fourth booster** | | | | | | | |
| | Pfizer bivalent (original and Omicron BA.4/BA.5) | 2 (8.0%) | 0 (0%) | 0 (0%) | 0 (0%) | 2 (40.0%) | 0 (0%) |
| | Pfizer monovalent XBB.1.5 | 6 (24.0%) | 0 (0%) | 3 (37.5%) | 2 (33.3%) | 0 | 1 (25.0%) |
| | Unboosted | 17 (68.0%) | 2 (100%) | 5 (62.5%) | 4 (66.7%) | 3 (60.0%) | 3 (75.0%) |
| **Fifth booster** | | | | | | | |
| | Moderna monovalent XBB.1.5 | 2 (8.0%) | 0 (0%) | 0 (0%) | 0 (0%) | 2 (40.0%) | 0 (0%) |
| | Unboosted | 23 (92.0%) | 2 (100%) | 8 (100%) | 6 (100%) | 3 (60.0%) | 4 (100%) |
| **Days between Fall 2023 vaccination and first serum sample collection** | | | | | | | |
| | Median (IQR) | 26 (20 - 32) | 23 (17 - 29) | 26 (19 - 33) | 24 (20 - 31) | 27 (26 - 36) | 21 (20 - 23) |
| **Days between Fall 2023 vaccination and second serum sample collection** | | | | | | | |
| | Median (IQR) | 201 (188 - 204) | 210 (204 - 217) | 194 (174 - 209) | 197 (187 - 229) | 201 (191 - 202) | 203 (192 - 203) |
| **Days between SARS-CoV-2 infection and first serum sample collection** | | | | | | | |
| | Median (IQR) | 570 (480 - 832) | - | 1095 (1095 - 1095) | 672 (651 - 674) | 551 (524 - 553) | 230 (99 - 351) |
| **Days between SARS-CoV-2 infection and second serum sample collection** | | | | | | | |
| | Median (IQR) | 729 (664 - 832) | - | 1247 (1247 - 1247) | 836 (817 - 874) | 708 (702 - 726) | 394 (254 - 530) |
| **Infecting variant*** | | | | | | | |
| | D614G | 1/16 (6.25%) | 0/2 (0%) | 1/8 (12.5%) | 0/6 (0%) | 0/5 (0%) | 0/4 (0%) |
| | BA.1 | 6/16 (37.5%) | 0/2 (0%) | 0/8 (0%) | 6/6 (100%) | 0/5 (0%) | 0/4 (0%) |
| | BA.2/BA.4 | 1/16 (6.25%) | 0/2 (0%) | 0/8 (0%) | 0/6 (0%) | 1/5 (20.0%) | 0/4 (0%) |
| | BA.4/5 | 4/16 (25.0%) | 0/2 (0%) | 0/8 (0%) | 0/6 (0%) | 4/5 (80.0%) | 0/4 (0%) |
| | XBB | 4/16 (25.0%) | 0/2 (0%) | 0/8 (0%) | 0/6 (0%) | 0/5 (0%) | 4/4 (100%) |

* Variants assigned based on date of infection

IQR, Interquartile range

**Table S4. Comparison of neutralization titers against XBB.1.5 variant after XBB.1.5 booster**

| Dunn's multiple comparisons test | 1 month | 6 months |
|---|---|---|
| D614G vs. XBB.1.5 | 0.06 | 0.0151 |
| B.1.617.2 vs. XBB.1.5 | 0.06 | 0.0135 |
| B.1.351 vs. XBB.1.5 | 0.69 | 0.1293 |
| BA.1 vs. XBB.1.5 | >0.9999 | >0.9999 |
| BA.4/5 vs. XBB.1.5 | >0.9999 | >0.9999 |
| XBB.1.5 vs. JN.1 | >0.9999 | >0.9999 |
| XBB.1.5 vs. KP.3 | 0.00 | 0.0008 |
| XBB.1.5 vs. XEC | 0.04 | 0.0665 |
| XBB.1.5 vs. KP.3.1.1 | <0.0001 | <0.0001 |
| XBB.1.5 vs. NB.1.8.1 | 0.00 | 0.0025 |
| XBB.1.5 vs. LP.8.1.1 | 0.01 | 0.0101 |
| XBB.1.5 vs. LF.7 | 0.00 | 0.0084 |
| XBB.1.5 vs. XFG | <0.0001 | <0.0001 |
| XBB.1.5 vs. XFC | 0.00 | 0.0435 |
| XBB.1.5 vs. BA.3.2.2 | >0.9999 | >0.9999 |
| XBB.1.5 vs. BA.3.2.1 | >0.9999 | >0.9999 |

**Table S5. Comparison of geometric mean neutralization titers for each variant at one month and six months**

| Wilcoxon matched-pairs signed rank test | P-Value |
|---|---|
| D614G 1M vs. D614G 6M | <0.0001 |
| B.1.617.2 1M vs. B.1.617.2 6M | <0.0001 |
| B.1.351 1M vs. B.1.351 6M | <0.0001 |
| BA.1 1M vs. BA.1 6M | <0.0001 |
| BA.4/5 1M vs. BA.4/5 6M | <0.0001 |
| XBB.1.5 1M vs. XBB.1.5 6M | <0.0001 |
| JN.1 1M vs. JN.1 6M | <0.0001 |
| KP.3 1M vs. KP.3 6M | <0.0001 |
| KP.3.1.1 1M vs. KP.3.1.1 6M | <0.0001 |
| XEC 1M vs. XEC 6M | <0.0001 |
| LP.8.1.1 1M vs. LP.8.1.1 6M | <0.0001 |
| LF.7 1M vs. LF.7 6M | <0.0001 |
| NB.1.8.1 1M vs. NB.1.8.1 6M | <0.0001 |
| XFG 1M vs. XFC 1M | <0.0001 |
| XFC 1M vs. XFC 6M | <0.0001 |
| BA.3.2.1 1M vs. BA.3.2.1 6M | <0.0001 |
| BA.3.2.2 1M vs. BA.3.2.2 6M | <0.0001 |
